# Asymbiotic nitrogen fixation is greater in soils under long-term no-till versus conventional tillage

**DOI:** 10.1101/502757

**Authors:** David W. Franzen, Patrick W. Inglett, Caley K. Gasch

## Abstract

A series of N-rate experiments was previously conducted in spring wheat, corn and sunflower in North Dakota indicated that less N was required when fields were in 6-years or more continuous no-till compared to conventional till. The objective of this study was to determine whether part of the reason for the decreased requirement for N was the greater activity of asymbiotic N-fixing organisms. Twelve paired-samplings were conducted in 2018. A surface 0-5cm deep sample was obtained in a long-term no-till field directly across the fence/road from a similar soil in conventional till. Samples were incubated in an acetylene-reduction procedure to estimate N fixation rate. Ten of twelve paired samplings had greater asymbiotic N fixation compared to the conventional till counterpart. This indicates that long-term no-till soils support greater N production from soil microorganisms than conventional till soils, which would result in lower input costs to no-till farmers.

## Introduction

Microbial biomass is usually greater in no-till soils than soils in conventional-till systems (Feng et al., 2003; Helgason et al., 2009). Analysis of over 100 nitrogen (N)-rate trials in spring wheat and durum wheat, over 120 N-rate trials in corn and over 30 N-rate trials in sunflower conducted in North Dakota shows that long-term no-till sites (greater than 6 years continuous no-till management) require less N to achieve maximum economic yield than conventionally tilled sites (Franzen, 2016; 2017; 2018).

It is possible that one explanation for the reduced N required in no-till sites is generally greater N use efficiency and protection from leaching/denitrification due to microbial N uptake. Another possible reason for reduced N requirement in long-term no-till is greater N fixation from asymbiotic N-fixing organisms. These are free-living organisms and do not have a symbiotic relationship with plant roots as do Rhizobium, Bradyrhizobium, or other symbiotic bacteria important to legumes. According to Roper and Gupta (2016), most soil nitrogenase enzyme is found in species within the Bacteria and Archaea domains. A limitation to the activity of asymbiotic N-fixing organisms is the food source. Asymbiotic N-fixing bacteria are present in nearly all cultivated soils (Wilson, 1958).

Long-term no-till fields generally have a greater supply of C and more diverse C sources compared to a conventional-tilled soil (Awale et al., 2013; Smith et al., 2016). Also, the greater stable aggregation in a no-till soil would provide a habitat for N-fixers and protect them from biocides that they might otherwise encounter (Gupta and Roper, 2010).

Microbial communities found in stable aggregates under no-till have a greater range of soil organic matter cycling-process rates than similar microbes in conventional systems where aggregates are disturbed or destroyed. No-till management systems are associated with greater number of N-fixing genes compared to soil from conventional-till fields (Smith et al., 2016).

There is no literature specifically comparing asymbiotic N-fixing in soils due to tillage differences. The objective of this study is to determine whether asymbiotic N-fixing is different in long-term no-till fields compared to adjacent fields in conventional tillage.

## Methods

Twelve paired sites were identified and sampled in North Dakota between April 26 and May 1, 2018. Each soil in a pair was within the same mapped, and confirmed through a field visit, soil series as its paired comparison. The NRCS (Natural Resource Conservation Service) Web Soil Survey was used as a guide to which area of the paired fields would represent a similar soil at each paired site (https://websoilsurvey.sc.egov.usda.gov/App/WebSoilSurvey.aspx). One soil of each pair was located in a no-till field with at least 6 years continuous no-till. No-till for the purposes of these comparisons included a ‘purist’ no-till, with only the seed disc cutting through the residue to the soil; strip-till, where a residue managers remove the residue from about 20% of the soil surface and a thin shank penetrates to the soil to about 20cm in depth, but leaves the rest of the soil undisturbed; and one-pass seeding, which is a shallow tillage using field cultivator shovels no deeper than 5cm at the time of seeding. The no-till location was directly across the fence, or road, no more than 50 meters away from the conventional tillage location.

Locations of the sites are provided in Figure 1. The locations were widely distributed within North Dakota. Samples were obtained during spring thaw before any spring field work was performed. The frost depth at the four northernmost sites was about 15cm below the soil surface, and frost depth at the remaining sites was about 30 cm below the surface. For each sample, the residue was brushed from the soil surface at a location about 40 meters inside the field boundary to avoid end-rows. A clean knife was used to scribe a 10 cm diameter cylinder of soil 5 cm deep. The cylinder was lifted from the soil using a stainless steel spatula and wrapped in aluminum foil to keep the soil intact as much as possible. The wrapped cylinder was then placed in a 11.4-cm diameter, 8.3-cm deep Ziploc^®^ screw-lid container and placed in a cooler. The knife and spatula were cleaned of soil between sites. Samples were stored in a refrigerator at -14°C until ready to ship for analysis. Previous crop and soil series for each site are listed in Table 1.

**Figure 1.**
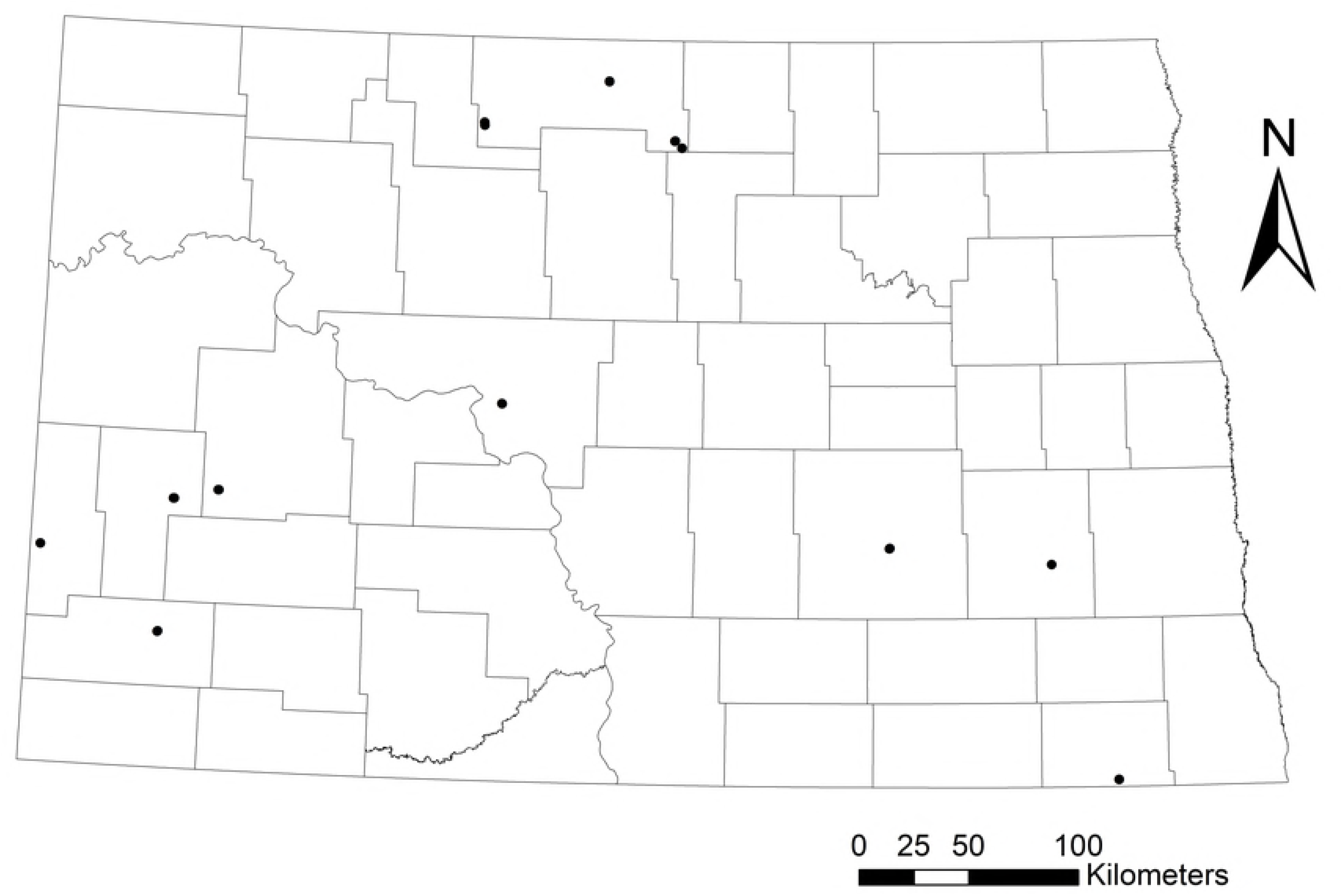
Locations of twelve paired-sample sites.

Samples were shipped to and analyzed by the Wetland Biogeochemistry Laboratory, University of Florida, Gainesville, FL, USA. Despite careful sampling, wrapping and shipping procedures, the samples did not arrive in a state that would allow an intact incubation. The soils were too dry at the time of sampling for them to remain as an undisturbed unit. The soils were therefore incubated separately as a bulk sample. Estimated N fixed per day utilized a 3:1 acetylene to N reduction ratio suitable for arable soils (Seitzinger and Garber, 1987).

**Table 1.** Selected site characteristics.

| Sample location | Tillage | Previous Crop | Soil Series |
| --- | --- | --- | --- |
| Amidon | Conventional | Spring wheat† | Rhodes sil‡, fine, smectitic, frigid, leptic Vertic Natrustolls |
|  | No-till | Spring wheat |  |
| Barton | Conventional | Corn | Embden fsl, coarse loamy, mixed, superactive, frigid, Typic Argiustolls |
|  | No-till | Corn |  |
| Beach | Conventional | Spring wheat | Chama sil, fine-silty, mixed, superactive, frigid, Typic Calciustolls |
|  | No-till | Corn |  |
| Belfield | Conventional | Spring wheat | Chama sil, fine-silty, mixed, superactive, frigid, Typic Calciustolls |
|  | No-till | Winter wheat |  |
| Bottineau | Conventional | Spring wheat | Overly sicl, fine-silty, mixed, superactive, frigid Pachic Hapludolls |
|  | No-till | Spring wheat |  |
| Dickinson | Conventional | Summer fallow | Morton sil, fine-silty, mixed, superactive, frigid Typic Argiustolls |
|  | No-till | Spring wheat |  |
| Jamestown | Conventional | Field pea/pea cover crop | Barnes l, fine-loamy, mixed, superactive, frigid Calcic Hapludolls |
|  | No-till | Soybean |  |
| Lansford | Conventional | Spring wheat | Hamlet l, fine-loamy, mixed, superactive, frigid Typic Calciaquolls |
|  | No-till | Spring wheat |  |
| Riverton | Conventional | Soybean | Williams l, fine-loamy, mixed, superactive, frigid, Typic Argiustolls |
|  | No-till | Corn |  |
| Rutland | Conventional | Soybean | Colvin sicl, fine-silty, mixed, superactive, frigid Pachic Hapludolls |
|  | No-till | Corn |  |
| Valley City | Conventional | Spring wheat | Hamerly l, fine-loamy, mixed, superactive, frigid Aeric Calciaquolls |
|  | No-till | Soybean |  |
| Willow City | Conventional | Corn | Hecla lfs, sandy, mixed, frigid Oxyaquic Hapludolls |
|  | No-till | Corn |  |

† Previous crop scientific names: Spring wheat/winter wheat – *Triticum aestivum*, L.; Field pea – *Pisum sativum*, L.; Soybean – *Glycine max* L., Merr.; Corn – *Zea mays*, L.
‡ Texture abbreviations: l – loam; sil – silt loam; sicl – silty clay loam; lfs – loamy fine sand; fsl – fine sandy loam

Nitrogen fixation potential of soil was assessed with the acetylene reduction assay using the procedures of Inglett (2013) and Matson et al. (2015). Briefly, soil samples were stored at 4° C until analysis (no more than 3 days). Soil samples were homogenized by hand mixing and added to 125 mL glass jars with metal, sealing lids which were ported to receive a luer connection, stopcock and syringe needle port. Two samples of field moist soil were prepared for each site with an additional sample taken for separate dry weight determination at 105° C.

To one set of prepared jars, acetylene (generated by adding water to CaC_2_) was added to each jar to enrich the headspace to approximately 10% by volume. Gas samples were taken at 4, 12, and 48 hours of incubation for determination of ethylene production from both acetylated and control (non-acetylated) soils. Ethylene values of acetylated jars were then corrected for ethylene in both the added acetylene (gas blanks) and that produced by the soil in the unacetylated controls. Gas samples were analyzed for ethylene using a Shimadzu GC-8A gas chromatograph equipped with a flame ionization detector (110°C) and a Poropak-N column (80°C). Two standard gases (1 ppm and 10 ppm; Scott Specialty Gases, Inc., Plumsteadville, PA) were used to calibrate the measurement, which was expressed as nmol C_2_H_4_ g dw^-1^ soil d^-1^. The data were subjected to statistical analysis in SAS 9.2 (SAS Institute, Cary, NC, USA) as a paired t-test, using a 95% confidence interval.

An estimate of N reduced using the acetylene reduction values was made. This estimate is based on several assumptions: the incubation procedure is similar to *in situ* processes and N fixation only occurs in the surface 5 cm of soil. One estimate of the N contribution from conventionally tilled soil to a wheat crop is about 10 kg N ha^-1^ per season (Kennedy and Islam, 2001).

## Results

Of the twelve paired sites, there were statistical differences (P<0.05) at ten sites (Table 2, Figure 1), where there was greater acetylene reduction in the no-till sample compared to the conventional-till sample. At three sites, there was no detectable acetylene reduction activity in the conventional-till site. The Lansford and Bottineau sites were not different in acetylene reduction compared to the conventional-till site. It is possible that what the no-till farmers considered to be ‘conventional tillage’ at these sites was actually a one-pass seeding. The higher rates of acetylene reduction in the conventional till soils at these sites compared to reduction rates at other conventional till sites over the study suggest that the true tillage management was mischaracterized.

**Table 2.** Soil nitrogenase activity of twelve paired samplings, no-till and conventional till.

| Sample location | Tillage | nmolC<sub>2</sub>H<sub>4</sub> reduced g<sup>-1</sup> DW soil day<sup>-1</sup> † | Estimated g N fixed ha<sup>-1</sup>day<sup>-1</sup> ‡ |
| --- | --- | --- | --- |
| Amidon | Conventional | Undetectable | 0 |
|  | No-till | 1.10 | 7.2 |
| Barton | Conventional | 0.36 | 2.4 |
|  | No-till | 1.12 | 7.3 |
| Beach | Conventional | Undetectable | 0 |
|  | No-till | 1.07 | 7.0 |
| Belfield | Conventional | 0.80 | 5.2 |
|  | No-till | 0.93 | 6.1 |
| Bottineau | Conventional | 0.77 | 5.0 |
|  | No-till | 0.75 | 4.9 |
| Dickinson | Conventional | 0.72 | 4.7 |
|  | No-till | 1.66 | 10.8 |
| Jamestown | Conventional | 0.71 | 4.6 |
|  | No-till | 1.15 | 7.5 |
| Lansford | Conventional | 0.48 | 3.1 |
|  | No-till | 0.49 | 3.2 |
| Riverton | Conventional | Undetectable | 0 |
|  | No-till | 1.28 | 8.4 |
| Rutland | Conventional | 0.83 | 5.4 |
|  | No-till | 1.38 | 9.0 |
| Valley City | Conventional | 0.72 | 4.6 |
|  | No-till | 2.02 | 13.2 |
| Willow City | Conventional | 0.35 | 2.3 |
|  | No-till | 0.59 | 3.9 |

†nanomoles C<sub>2</sub>H<sub>4</sub> reduced gram<sup>-1</sup> of dry weight soil per day
‡LSD 5% between No-till and Conventional-till reduction is 0.1 nmolC<sub>2</sub>H<sub>4</sub> reduced g<sup>-1</sup> DW soil day<sup>-1</sup>.
¶Assumes 1.4 g soil cm<sup>-3</sup>, ratio of C<sub>2</sub>H<sub>4</sub> : N<sub>2</sub> reduced is 3:1. Reduction under incubation conditions.

The previous crop is noted in Table 1 for the record, but it is unlikely that a previous crop had any influence on acetylene reduction results. For example, 3 sites had spring wheat as previous crop before no-till and conventional till soils (Amidon, Lansford and Bottineau). Although the acetylene reduction rates for conventional and no-till soils at Lansford and Bottineau soils were not different, the acetylene reduction rates for soils at Amidon were greatly different.

An estimate of g N fixed ha^-1^ day^-1^ is provided (Table 2). If we use the mean difference in these comparisons of 0.48 nmol C_2_H_4_ reduced g^-1^ DW soil day^-1^ in conventional-till compared with 1.13 nmol C_2_H_4_ reduced g^-1^ DW soil day^-1^ in no-till, and a factor of 2.35 for C_2_H_4_ to N, we can calculate an estimate of the growing season N contribution in no-till. That value would be 23.5 kg N ha^-1^ for a 100 day growing season^-1^; an increase of 13.5 kg N ha^-1^ per season over a conventional till soil. The 13.5 kg N ha^-1^ increase available in no-till from conventional till is approximately one-quarter to one-third of the N of the N reduction indicated from N-rate studies on no-till compared to conventional till in North Dakota.

## Discussion

Although several studies have indicated that greater asymbiotic N fixation is possible under no-till (Gupta and Roper, 2010; Smith et al., 2016) this is the first study to indicate that greater asymbiotic N is produced in soils under long-term no-till management. This study will hopefully lead to more careful examination of paired fields, or carefully constructed tillage experiments. It takes time to establish credible no-till experimental sites. Establishing credible tillage experiments are difficult for most researchers. Experimental farms often have space allotments that reduce the area that could be devoted to tillage research. Sufficient space must be allocated so that no-till treatments are not driven on to reach a tillage treatment. It is essential that timing of planting be conducted when the soil is fit for the tillage practice. However, in most tillage studies, a choice has to be made whether to plant when the conventional till plots are fit, or the no-till plots are fit. A farmer would choose the ideal planting date for their tillage system. A researcher usually gives up something because they cannot go back and forth multiple times to plant on the ideal date for each tillage treatment. We therefore have chosen the paired-sample method between closely neighboring fields with similar soils to take advantage of the farmer ability to conduct their operations in the timeliest manner, whether the sites were no-till or conventional till.

This study examined only one 0-5 cm deep sample within a no-till location. There is nothing in this study to indicate the spatial variability that might be present within 1m, 5m, 100m which would indicate the uniformity or variability of N fixation in the field. Further studies are required to provide a range of N fixation values for consideration if an N credit towards N rate reduction is to be constructed or modified in the future.

## Conclusions

Long-term no-till soils tended to have greater asymbiotic N fixation than neighboring soils under conventional tillage. This indicates that reduced fertilizer N requirements for corn, wheat and sunflower in North Dakota long-term no-till fertilizer recommendations have at least one explanation. Future studies should include the spatial nature of asymbiotic N fixation in no-till fields and whether differences are present in no-till variations, including penetration of soil with the seeding disc only, or strip-till, or shallow (<5cm deep) tillage in one-pass seeding systems, which are all generally categorized as ‘no-till’. This study also provides evidence that greater farm sustainability is possible from transitioning to no-till, due to reduced N inputs required to produce a crop.

